# Development and optimization of biologically contained Marburg virus for high-throughput antiviral screening

**DOI:** 10.1101/2022.07.29.502006

**Authors:** Bert Vanmechelen, Joren Stroobants, Winston Chiu, Lieve Naesens, Joost Schepers, Kurt Vermeire, Piet Maes

**Affiliations:** Department of Microbiology, Immunology and Transplantation, Rega Institute, Laboratory of Clinical and Epidemiological Virology, KU Leuven, Herestraat 49 – box 1040, 3000 Leuven, Belgium; Department of Microbiology, Immunology and Transplantation, Rega Institute, Laboratory of Virology and Chemotherapy, KU Leuven, Herestraat 49 – box 1043, 3000 Leuven, Belgium

## Abstract

Comparable to the related Ebola virus, Marburg virus is an emerging zoonotic pathogen that causes hemorrhagic fever with a high mortality rate. Therefore, handling of Ebola virus and Marburg virus is limited to biosafety level 4 facilities, of which only a limited number exists worldwide. However, researchers have developed several virus alternatives that are safe to handle in lower biosafety settings. One particularly interesting approach is the engineering of biologically contained Ebola virus by removing an essential gene from the virus genome and providing this missing gene in trans in a specific cell line. Because the virus is confined to this specific cell line, this results in a system that is safe to handle. So far, Ebola virus is the only virus for which biological containment has been reported. Here, we describe the first successful rescue of biologically contained Marburg virus and demonstrate that biological containment is also feasible for other filoviruses. Specifically, we describe the development of containment cell lines for Marburg virus through lentiviral transduction and show the growth and safety characteristics of eGFP-expressing, biologically contained Marburg virus in these cell lines. Additionally, we exploited this newly established Marburg virus system to screen over 500 compounds from available libraries. Lastly, we also validated the applicability of our biologically contained Marburg virus system in a 384-well format, to further illustrate the usefulness of this novel system as an alternative for high-throughput MARV screening of compound libraries.

## Introduction

EBOV (species *Zaire ebolavirus*) and MARV (species *Marburg marburgvirus*) are the most prevalent members of the genera *Ebolavirus* and *Marburgvirus*, respectively [1]. Since their discovery more than 40 years ago, both viruses have caused several outbreaks of viral hemorrhagic fever, all characterized by a high mortality rate [2]. Although research in recent years has led to the development of effective EBOV vaccines, there is no licensed MARV equivalent currently available [3, 4, 5]. In addition, MARV is also relatively understudied when it comes to treatment options, and there are no licensed therapeutics to treat Marburg virus disease (MVD). For Ebola virus disease (EVD), a recent clinical trial showed that two monoclonal antibodies, MAb114 (Ebanga) and REGN-EB3 (Inmazeb), improved disease outcome, but the overall mortality remained high [6]. Because of this lack of suitable preventive measures and treatment options, in combination with the high risk these viruses pose for the personnel handling them, both EBOV and MARV are classified as BSL-4 agents [7]. As the number of BSL-4 facilities worldwide is limited, this classification has significantly hindered the advancement of filovirus research. In the last few decades, several virus alternatives have been developed that allow filovirus research to be conducted in lower biosafety settings, but each of these systems has its own limitations, often necessitating the validation of obtained results with replication-competent filovirus [8, 9].

In 2008, Peter Halfmann and colleagues showed that it is possible to confine EBOV to a specific cell line by removing one of the essential genes (VP30) from the virus genome and providing the missing protein in trans in the cell line of choice [10]. This results in the production of EBOV that is phenotypically near indistinguishable from wild-type virus, but which is safe to handle. Although the functioning of VP30 has not been fully resolved, it is known to act as an enhancer of virus transcription and it has been shown to be indispensable for virus replication [11, 12]. Based on this design, we have previously shown that it is possible to perform high-throughput screening of antiviral compounds using biologically contained EBOV, by creating suitable containment cell lines in which EBOV VP30 is stably integrated into the host genome via lentiviral transduction [13]. In this study, we show that through a series of modifications the rescue of biologically contained virus is also possible for MARV. We describe here the characterization of this biologically contained MARV and optimize its use in 96- and 384-well formats. Lastly, we also validated the usability of our novel assay for antiviral compound screening using two repurposing compound libraries, the Pandemic Response Box (400 compounds) and the Covid box (160 compounds), made available by the Medicines for Malaria Venture (MMV) [14, 15]. All 560 compounds in these two libraries are known drugs or drug-like compounds that have proven or predicted activity against bacteria, viruses or fungi, including the known MARV inhibitors remdesivir and apilimod [16, 17].

## Materials and methods

### Cell lines

Cell lines used for this study were native hamster kidney fibroblasts (BHK-21; BHK-21(C-13), ATCC, Manassas, VA, USA) and a derived cell line constitutively expressing T7 polymerase (BSR-T7/5; kindly provided by Dr. K. Conzelmann, Friedrich-Loeffler-Institut, Germany), African green monkey kidney cells (Vero E6; Vero C1008, ATCC) and human hepatocellular carcinoma cells (Huh-7; kindly provided by Ralf Bartenschlager, University of Heidelberg, Germany) [18]. BHK-21 cells were maintained in minimum Essential Medium (MEM) REGA-3 (Thermo Fisher Scientific, Waltham, MA, USA) and all other cell lines in Dulbecco’s Modified Eagle Medium (DMEM; Thermo Fisher Scientific). All media was supplemented with 10% fetal bovine serum (FBS; Biowest, Nuaillé, France) 1% sodium bicarbonate (Thermo Fisher Scientific) and 200mM L-glutamine (Thermo Fisher Scientific). Additionally, 1% Non-Essential Amino Acids (Thermo Fisher Scientific) was added for Huh-7 cells. 1% Penicillin-Streptomycin (Thermo Fisher Scientific) was used as antibiotic for each cell line, supplemented with 0.5% Geneticin (InvivoGen, San Diego, CA, USA) for BSR-T7/5 cells, and 1% Gentamicin (Thermo Fisher Scientific) and 0.2% Fungizone (Thermo Fisher Scientific) for Vero E6 cells. Additional antibiotics and modified serum concentrations were used after lentiviral transduction and during assays (see below).

### Plasmids

pCAGGS plasmids encoding MARV L, NP, VP30 and VP35, as well as the T7-3M-Luc-5M minigenome plasmids, based on the Musoke strain, were kindly provided by Prof. Stephan Becker [19]. Codon-optimized variants of the MARV pCAGGS helper plasmids were generated from pUC57 plasmids containing the full-length codon-optimized sequences, synthesized and codon-optimized by GenScript Biotech (GenScript Biotech, Piscataway, NJ, USA). pUC57 vectors containing the codon-optimized sequences of VP30, VP35, NP and L, and the pCAGGS vector were digested with EcoRI-HF and NheI-HF (New England Biolabs, Ipswich, MA, USA) and ligated with the Quick Ligation kit (New England Biolabs).

A plasmid encoding an eGFP-containing MARV antigenome was generated through assembly of fragments using the NEBuilder HiFi DNA assembly cloning kit (New England Biolabs). The vector backbone, T7 promoter, virus leader, virus trailer, HdVRz and T7 terminator sequences were derived from the T7-3M-Luc-5M vector. Fragments covering the NP and L genes were derived from the corresponding pCAGGS helper plasmids. The rest of the antigenome, from the intergenic region in front of the VP35 gene to the intergenic region behind the VP24 gene, with an eGFP gene replacing the VP30 coding region, was synthesized based on MARV strain Musoke (GenBank: DQ217792) by GenScript Biotech, in a pUC57 vector. This fragment was digested with NotI-HF and SmaI (New England Biolabs), while all other fragments were amplified by PCR using the Q5 HotStart High-Fidelity 2X master mix (New England Biolabs). All plasmids were sequence-verified with Sanger sequencing (Macrogen Europe, Amsterdam, The Netherlands) before use.

### Lentivirus production and transduction

A lentiviral construct containing MARV VP30 linked to a blasticidin S-deaminase (BSD) resistance gene by means of an internal ribosomal entry site (IRES) cassette was used to create stably transduced cell lines expressing VP30. The MARV lentivirus was made by modifying the existing EBOV VP30 constructs previously described in [13]. Assembly of PCR fragments with the NEBuilder HiFi DNA assembly cloning kit (New England Biolabs) was used to replace the EBOV VP30 by a (codon-optimized) MARV VP30 gene. A vector containing the 14 Amino-acid Simian Virus 5 epitope Pk (V5) in front of the VP30 gene was generated in the same way.

A day prior to transfection, HEK293FT cells (Thermo Fisher Scientific) were seeded in T-25 flasks at a cell density of 3 × 10^6^ cells in DMEM (Thermo Fisher Scientific), supplemented with 10% FBS (Biowest), to achieve a confluency of 50-70% on the day of transfection. Transfection was performed with Lipofectamine LTX & PLUS Reagent (Thermo Fisher Scientific). LTX solution and transfection mixes containing 3 μg of lentiviral MARV vector, 5.83 μg of psPAX2 vector, 3.17 μg of pMD2.G vector and 12 μL PLUS reagent were prepared in serum-free Opti-MEM (Thermo Fisher Scientific). Following a five-minute incubation at room temperature, solutions were mixed and incubated for an additional 20 minutes. The medium from the T-25 flasks was replaced with 5 mL fresh growth medium, after which transfection complexes were added. Cells were then incubated at 37°C for 21 hours. Sodium butyrate was added to a final concentration of 10 mM and cells were incubated for 3 hours, after which the medium was replaced with 5 mL fresh growth medium. Virus-containing supernatants were harvested into 15 mL conical tubes 24 hours after sodium butyrate addition and centrifuged at 2,000 g for 15 minutes at 4°C to pellet cell debris. Aliquots of viral supernatants were stored at −80°C until they were used for the transduction of cell lines, which was done according to the ViraPower HiPerform T-Rex Gateway Expression System (Thermo Fisher Scientific) manufacturer’s protocol. Six μg/ml Polybrene (Sigma-Aldrich, Saint-Louis, MO, USA) was used to increase transduction efficiency. Following transduction, cell medium was supplemented with 10-40 μg/ml blasticidin (InvivoGen) during passaging.

### MARV rescue

Vero E6 cells transduced with codon-optimized MARV VP30 (Vero E6-MARV-CO-VP30) were seeded in a 6-well plate (800.000 cells/well). Following overnight incubation, cells were transfected with 500 ng MARV antigenome, 500 ng T7 polymerase, 500 ng pCAGGS-MARV-CO-NP, 1000 ng pCAGGS-MARV-CO-L, and 250 ng pCAGGS-MARV-CO-VP35, using 6:1 TransIT-LT1 Transfection Reagent (Mirus Bio, Madison, WI, USA). One and six days after transfection, the medium was replaced by fresh 2% FBS medium. Thirteen days post-transfection, 1 ml supernatant from wells showing eGFP expression was collected, centrifuged at 17,000 g for three minutes and transferred to a T-75 flask filled with Vero E6-MARV-CO-VP30 cells, seeded one day prior. After seven days, supernatant was collected and used to infect additional T-75 flasks of Vero E6-MARV-CO-VP30 cells, from which, after seven days, the supernatant was collected, centrifuged at 17,000 g for three minutes and subsequently aliquoted and stored at −80°C.

### Western blotting

Cells were lysed in ice-cold Nonidet P-40 buffer [50 mM Tris·HCl (pH 8.0), 150 mM NaCl, 1% Nonidet P-40] (Thermo Fisher Scientific) supplemented with cOmplete protease inhibitor cocktail (Roche, Basel, Switzerland) and phenylmethylsulphonyl fluoride (Sigma-Aldrich). Lysates were centrifuged at 17,000 g for 10 min to pellet nuclei and debris. For sodium dodecyl sulfate (SDS) gel electrophoresis, samples were boiled in reducing 2X Laemmli sample buffer (120 mM Tris·HCl (pH 6.8), 4% SDS, 20% glycerol, 100 mM dithiothreitol, 0.02% bromophenol blue). Equal volumes of sample were separated on 4–12% or 10% Criterion XT Bis-Tris Precast gels (Bio-Rad, Hercules, CA, USA) using XT MES Running buffer (Bio-Rad). After electrophoresis, gels were blotted onto polyvinylidene difluoride membranes (Bio-Rad) using the Trans-Blot Turbo transfer system (Bio-Rad). Membranes were blocked for one hour with 5% nonfat dried milk in TBS-T (20 mM Tris·HCl (pH 7.6), 137 mM NaCl, 0.05% Tween-20). After one hour of incubation with primary antibodies, membranes were washed and incubated with secondary horseradish peroxidase-labeled antibody. SuperSignal West Femto chemiluminescence reagent (Thermo Fisher Scientific) was used for detection in conjunction with a ChemiDoc MP system (Bio-Rad). THE V5 Tag Antibody (1:2000, GenScript Biotech) was used as primary antibody for samples and mouse anti-clathrin heavy chain clone 23 (1:2,000; BD Biosciences, Franklin Lakes, NJ, USA) for the detection of clathrin heavy chain as a loading control. Goat anti-mouse HRP (1:10,000; Agilent, Santa Clara, CA, USA) was used as secondary antibody.

### Virus titration

Virus stock titers were determined by ‘plaque’ assay. Vero E6-MARV-CO-VP30 cell lines were seeded in 6-well plates and a ten-fold dilution series, covering ten dilutions (1×10^-1 - 1×10^-10) was made for each virus. Once confluent, cell medium was removed and 200 μl virus dilution was added to each well, using duplicate repeats for each concentration. Plates were kept in an incubator (37°C, 5% CO2), gently swirling the plates every 15 minutes. After one hour, 3 ml freshly prepared agarose-medium was added to each well. Agarose-medium was made by autoclaving a dilution of 17.6 μg/ml SeaKem ME agarose (Lonza, Basel, Switzerland) and heating it to 65°C. Once heated, the agarose was added to preheated (37°C) 2X Basal Medium Eagle without Earle’s salts (Thermo Fisher Scientific), supplemented with 10% FBS (Biowest), 200 mM L-glutamine, 1% NEAA, 1% Penicillin-Streptomycin, 1% Gentamicin and 0.2% Fungizone (all Thermo Fisher Scientific), in a 1:2 ratio. After cooling down to room temperature, plates were moved to an incubator for five days. Read-out was performed by counting the amount of eGFP+-cell clusters.

### RNA extraction and RT-qPCR

RNA was extracted from 100 μl of virus stock using a KingFisher Flex (Thermo Fisher Scientific) in combination with the MagMax Viral Pathogen kit II (Thermo Fisher Scientific), according to the manufacturer’s instructions. Reverse transcription quantitative PCR (RT-qPCR) assays were performed using the Quantstudio 5 Real-time PCR system (Applied Biosystems, Waltham, MA, USA). Premixes were prepared for each amplification reaction using TaqMan Fast Virus 1-Step Master Mix (Thermo Fisher Scientific), according to the manufacturer’s protocol. Primers and double-quenched FAM probes (Supplementary Table S1) were obtained from Integrated DNA technologies (Leuven, Belgium) and checked for specificity via BLAST (https://blast.ncbi.nlm.nih.gov/Blast.cgi). Primers and probes were designed based on the L gene sequences of Marburg virus strain Musoke (Genbank accession no. DQ217792) with an amplicon length of 125 base pairs. Standard curve DNA fragments were derived from PCR products of both L genes, generated with primers flanking the qPCR amplicon region. PCR products were purified with the Gel and PCR cleanup kit (Macherey-Nagel, Dueren, Germany) according to the manufacturer’s protocol. Five μL sample was used for each reaction, in a total reaction volume of 20 μL. Final concentrations of 500 nM of each primer and 250 nM probe were used per reaction. PCR run conditions were 1 cycle of 50°C for 5 min and 95°C for 20 sec, followed by 40 cycles of 95°C for 15 sec and 55°C for 60 sec. Data analysis was performed with the Quantstudio 3&5 Design and analysis software (Applied Biosystems, v1.5.1). Cq values were obtained via automated threshold determination. Standard curve runs were included in duplicate on each plate, and efficiencies were calculated based on the slope. qPCR runs with standard curve efficiencies not ranging between 90% and 110% or no-template controls showing amplification were excluded from the data.

### Nanopore sequencing

RNA was converted to cDNA and amplified by Sequence-Independent Single Primer Amplification as described by Goya et al. [20]. The resulting cDNA was prepared for Nanopore sequencing using the SQK-LSK110 kit (Oxford Nanopore Technologies (ONT), Oxford, UK) with the EXP-NBD114 barcoding expansion (ONT). The resulting library was loaded on a R9.4.1 flow cell and run on a GridION. Basecalling and barcode demultiplexing was done using the ont-guppy-for-gridion v4.2.3. The resulting reads were mapped against the plasmid design used for generation of the antigenome construct using Minimap2 v2.17-r941, followed by Medaka v1.0.1 for consensus polishing and variant calling [21].

### Compound screening

Compounds were freely provided to us by the Medicines for Malaria Venture in the form of DMSO solutions of 2 or 10 mM, pre-spotted on 96-well plates. For the initial screening of the two libraries, Vero E6-MARV-VP30-CO cells were seeded in 96-well plates (20,000 cells/well) and incubated at 37°C for 24 hours. The next day, an intermediate four-fold dilution series was made of each compound, after which 50 μl of each dilution was transferred to the cell-filled plate, to achieve a final compound concentration of 50, 12.5, 6.13 or 1.56 μM. Following a 1-hour incubation at 37°C, 200 PFU MARV-ΔVP30-eGFP was added to each well and the plates were returned to the incubator. After six days, high-content imaging was performed on an Arrayscan XTI (Thermo Fisher Scientific) as described previously [22]. For all plates, 6 μM Hoechst 33342 nucleic acid stain (Thermo Fisher Scientific) was added to each well at least 1 hour before images were taken, to serve as a nuclear counterstain for autofocusing and cell counting. HCS Studio (Thermo Fisher Scientific), using in-house developed protocols, was used to calculate the number and percentage of eGFP-positive cells. Further data analysis and graph plotting was done using GraphPad Prism v8.2.0.

For hit confirmation, additional compound was acquired through Evotec (Hamburg, Germany) and used to create fresh DMSO stocks. A two-fold dilution series over a wider concentration range (≥8 concentrations, starting at 50 μM) was used to assess more accurately the activity and cytotoxicity of each compound. Each dilution series was tested in triplicate and each assay was repeated at least twice to account for inter- and intra-experimental variability. Other assay parameters were identical to the initial screen and all plate handling and imaging procedures were performed as described above. The activity of confirmed hits was also determined in Huh-7-MARV-CO-VP30 cells. For this assay, 10,000 cells and 2,000 PFU MARV-ΔVP30-eGFP were used per well, and assay read-out was performed four days post-infection.

## Results

### Generation of VP30-expressing cell lines

Although the functioning of VP30 has not been fully resolved, it is known to act as an enhancer of virus transcription and it has been shown to be indispensable for virus replication [11, 12]. To create cell lines continuously expressing MARV VP30, a construct containing the VP30 gene with or without a V5-tag on the N-terminal side, coupled to a blasticidin-resistance gene by means of an IRES, was stably integrated into the host cell’s genome through lentiviral transduction (Figure 1A). To increase the chance of virus rescue, four different cell lines that had previously been successfully used for the rescue of filoviruses were selected for VP30 transduction: BSR-T7/5, BHK-21, Vero E6 and Huh-7 cells [23, 24, 25, 26, 27, 28]. Despite strong resistance of the cells against the used selection marker (blasticidin), VP30 expression appeared limited, especially in the Huh-7 and Vero E6 cells (Supplementary Figure 1). Initial attempts to improve MARV VP30 expression by using different resistance markers or higher antibiotic concentrations were unsuccessful. Because we hypothesized that the limited MARV VP30 expression could be attributed to low translation efficiency, a codon-optimized version of the VP30 gene was incorporated into the lentivirus design. Aside from the BHK-21 cells, which already showed the highest level of VP30 expression prior to codon optimization, all cell lines showed a clear improvement in VP30 expression following transduction with this codon-optimized construct.

**Figure 1:**
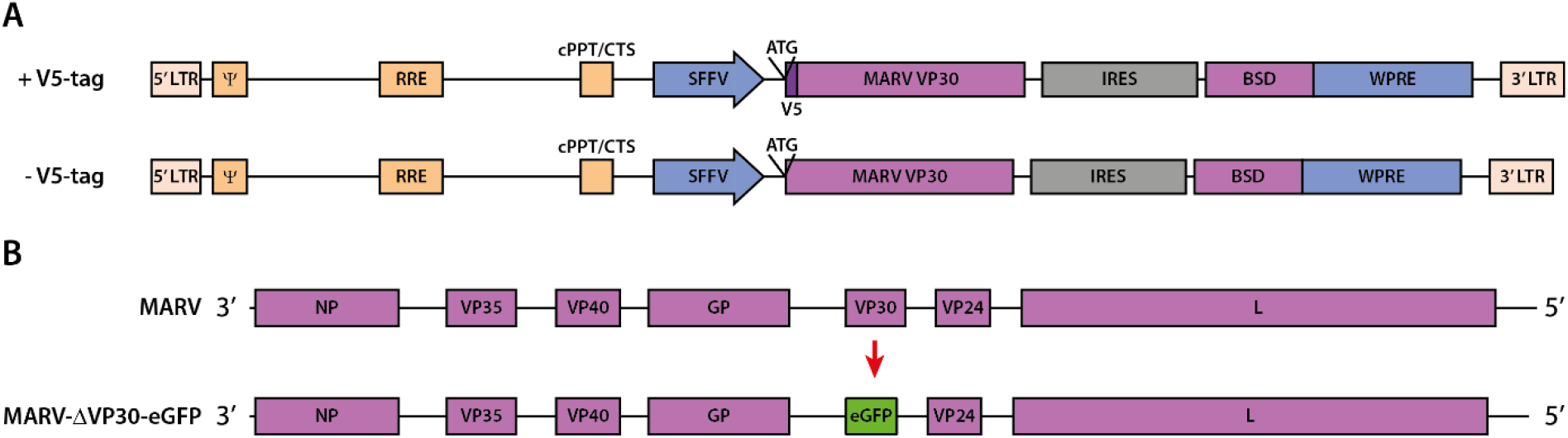
Schematic overview of the virus constructs used. (A) Lentiviral construct used for transduction, incorporating a (codon-optimized) MARV VP30 gene (with or without V5-tag) joined to a blasticidin resistance gene (BSD) using an internal ribosomal entry site (IRES). LTR = Long terminal repeat, RRE = Rev Response Element, cPPT/CTS = Central polypurine tract/central termination sequence, SFFV = Spleen focus-forming virus promoter, WPRE = Woodchuck hepatitis virus post-transcriptional regulatory element. (B) To create MARV-ΔVP30-eGFP, the ORF containing the VP30 gene is replaced by that of eGFP.

### Rescue of biologically contained MARV

A VP30-deficient MARV antigenome construct was made by replacing the VP30 open reading frame by that of eGFP (Figure 1B). Initial attempts to rescue virus from this construct using cells transduced with non-codon optimized MARV VP30 did not result in virus rescue, regardless of the cell line used, presumably due to insufficient availability of VP30. Furthermore, even though codon optimization improved VP30 expression in the transduced cell lines, attempts to rescue MARV through transfection of Huh-7-MARV-CO-VP30, BHK-21-MARV-CO-VP30 or BSR-T7/5-MARV-CO-VP30 cells and subsequent supernatant transfer to Vero E6-MARV-CO-VP30 cells were unsuccessful. Because codon optimization seemed necessary to reach sufficient VP30 expression, we hypothesized that the failure to rescue MARV could be attributed to an inadequate balance or lack of support proteins (VP35, NP and L). Therefore, all support plasmids were also codon-optimized and different plasmid ratios were evaluated, including the condition previously used for the rescue of biologically contained EBOV by our group, the conditions for MARV rescue previously described by Albariño et al. and Enterlein et al., as well as the balances used in the iVLP system of Wenigenrath et al., with or without the structural support plasmids [12, 13, 19, 26]. These conditions were evaluated in Huh-7-MARV-CO-VP30, BHK-21-MARV-CO-VP30 and BSR-T7/5-MARV-CO-VP30 cells, using both genomic and antigenomic MARV constructs, totaling more than 100 individual rescue attempts, which were all unsuccessful. Because even previously published rescue protocols did not result in virus growth, we hypothesized that the amount of VP30 being produced by the cells was still insufficient, or that the cells underwent cell death too fast to allow successful virus rescue, a phenomenon previously observed for BSR-T7/5 cells by Albariño et al. [26]. To address these issues, we decided to attempt virus rescue directly in Vero E6 cells. Vero E6-MARV-CO-VP30 and Vero E6-MARV-VP30 cells were transfected in 6-well format, using twice the amount of transfection reagent (6:1) and the plasmid ratios used by us for the rescue of biologically contained EBOV or those described by Albariño et al. (three replicates each) [13, 26]. On day eight post-transfection, respectively one and five cluster(s) (~20 cells) of eGFP-positive cells appeared in two replicates of the Vero E6-MARV-CO-VP30 cells transfected with the conditions described by Albariño et al. [26]. After an additional four days, cell medium from these two wells was used to infect fresh Vero E6-MARV-CO-VP30 cells, resulting in widespread eGFP expression and thereby confirming virus rescue.

### Virus characteristics and biological confinement

Nanopore sequencing was used to confirm the complete genome sequence of the two rescued viruses. An overview of the observed mutations for the two isolates is shown in Table 1. Isolate A was found to have two silent mutations in the L gene and an insertion of one additional adenosine residue in the polyadenylation signal following the VP30, or in this case eGFP, gene (5A>6A). Isolate B had only a single substitution in the non-coding region following the VP24 gene. To study the characteristics of both isolates, we first determined the concentration of infectious particles in our virus stocks by plaque assay, as well as the minimal virus dose needed to achieve maximal eGFP expression in transduced Vero E6 cells. For both MARV-ΔVP30-eGFP isolates, a loading dose of 100 plaque forming units (PFU)/well, corresponding to 0.005 PFU/cell, was found to consistently yield >90% eGFP-positive cells six days post-infection in all replicates, while lower doses failed to uniformly infect all cells or replicates (Figure 2). To evaluate the growth kinetics in greater detail, virus growth on four differently transduced cell lines was observed daily over a period of six days, using a low (0.01 PFU/cell) and a high (0.1 PFU/cell) viral loading dose. In cells expressing codon-optimized MARV VP30, initial eGFP expression could be observed already after one day, increasing rapidly until day four or six, depending on the used dose. Maximum eGFP expression, corresponding to ~95% of all cells being positive, was achieved two days faster when higher titers were used, with no notable difference between the two virus isolates (Figure 3), as also confirmed by RT-qPCR (Supplementary Figure S2). Conversely, when using non-transduced Vero E6 cells, no eGFP expression was observed after six days, confirming the confinement of all viruses to their respective cell lines. Interestingly, codon optimization of MARV VP30 was necessary for the growth of MARV-ΔVP30-eGFP. In Vero E6 cells transduced with a regular MARV VP30-construct, overall eGFP expression levels did not rise above the background noise observed in non-infected control wells, although occasionally small clusters of eGFP-expressing cells could be observed, indicating that virus growth is possible in this cell line but only limitedly (Supplementary Figure S3). We previously also established a cell line expressing EBOV VP30 [13]. Of note, when infecting these cells with MARV-ΔVP30-eGFP, equally distributed eGFP expression could be observed in all replicates, albeit at a very weak efficiency, making it difficult to distinguish eGFP-positive cells correctly from background noise (Supplementary Figure S3).

**Figure 2:**
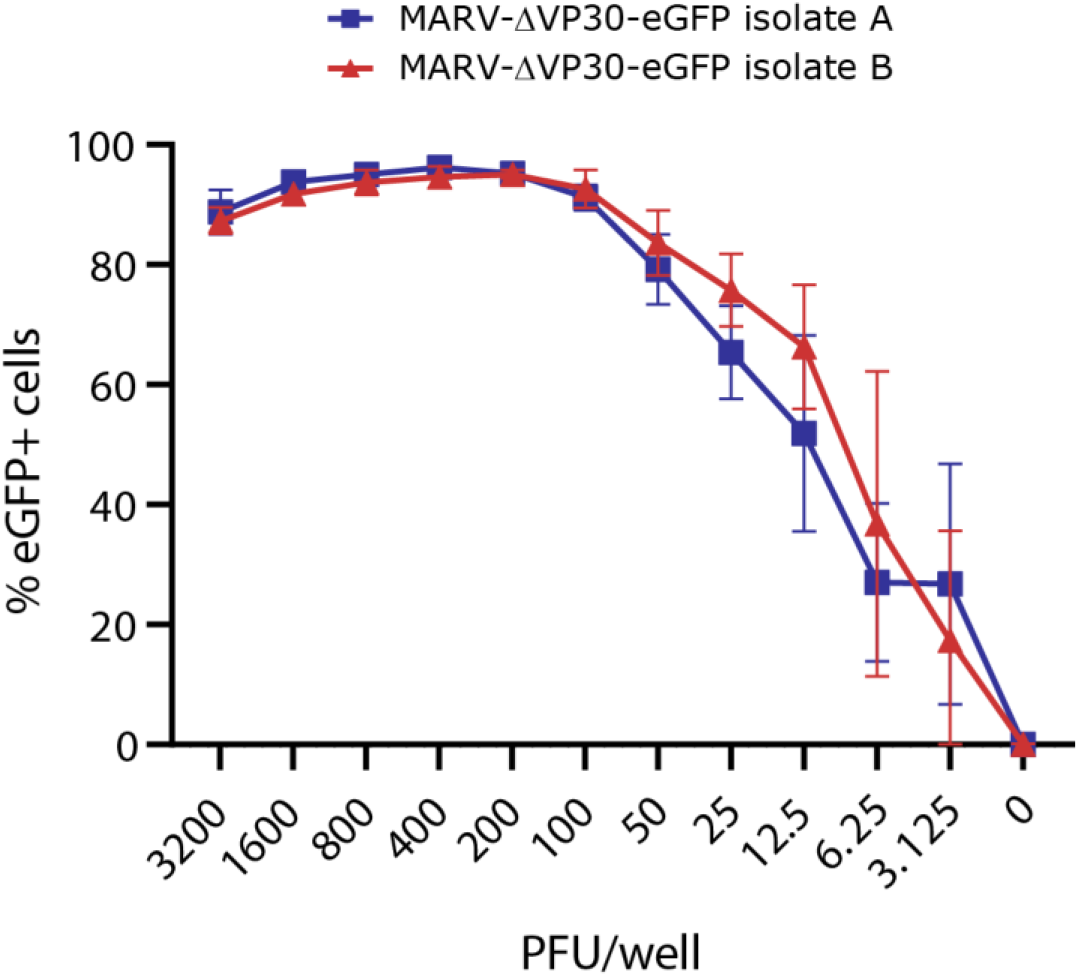
Minimal virus titer required for homogenous infection in 96-well plates. Vero E6-MARV-CO-VP30 cells were infected with MARV-ΔVP30-eGFP isolate A or B, respectively, and the fraction of eGFP-positive cells was determined by high-content imaging six days post-infection. Different viral titers tested are expressed in plaque forming units (PFU)/well. Eight replicates across two separate plates were performed per condition. Error bars denote standard deviation.

**Figure 3:**
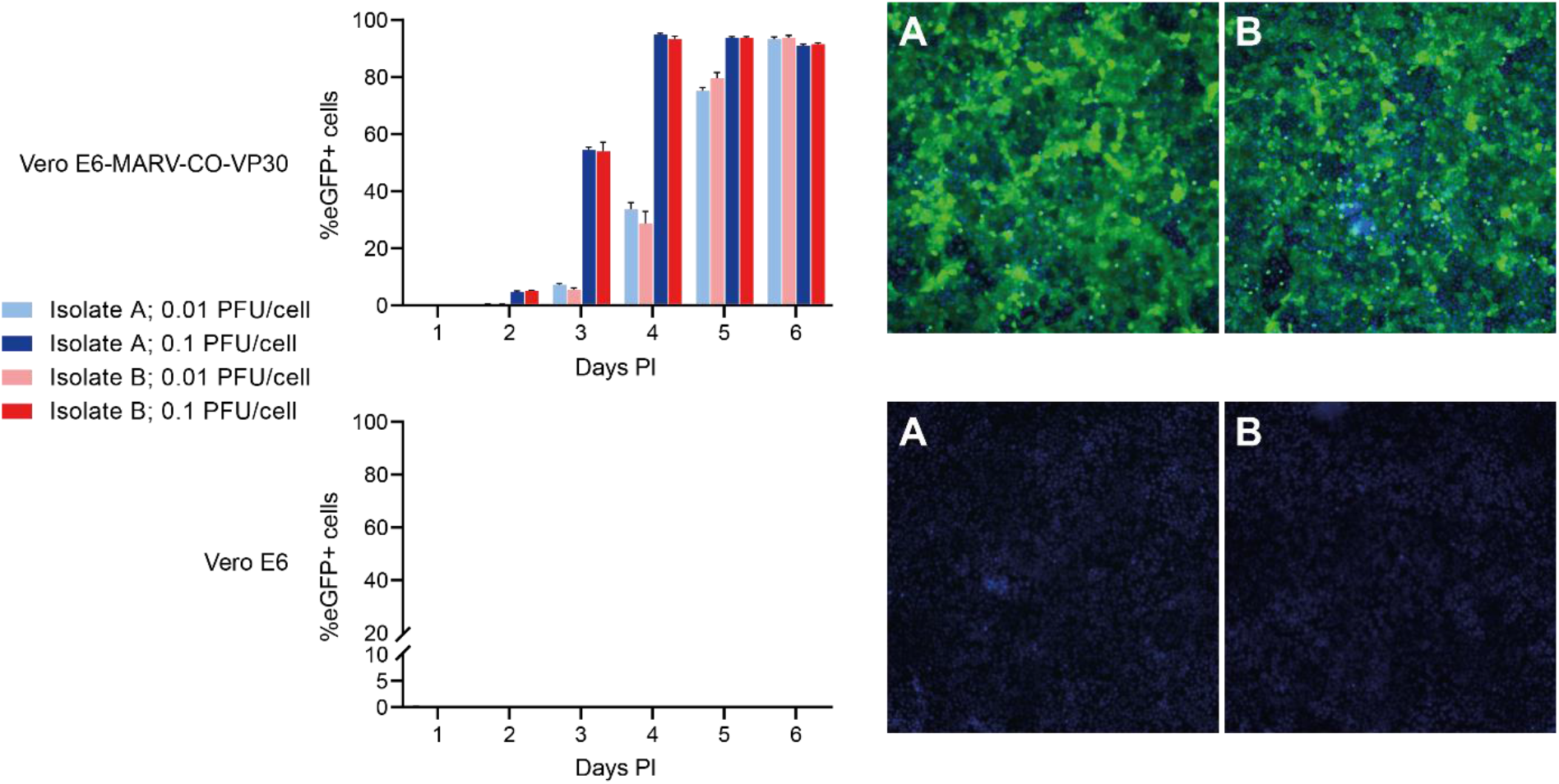
Growth kinetics and biological containment of MARV-ΔVP30-eGFP. The left graph shows the background-corrected fraction of eGFP-positive cells, measured by high-content imaging. The right column shows a representative image of a well infected with 0.1 PFU/well, five days post-infection. Green cells express eGFP and all cells are nuclear stained with Hoechst 33342 (blue). PFU = plaque forming units. At least three replicates are included for each condition. Error bars denote standard error.

**Table 1:** Mutation overview of the rescued virus isolates

| Virus isolate | Mutation (nt)* |
| --- | --- |
| MARV-ΔVP30-eGFP A | G9885GA [G10011GA] |
|  | A12440G [A12566G] |
|  | A12467G [A12593G] |
| MARV-ΔVP30-eGFP B | T10971A [T11097A] |

\*Position of each mutation in the virus isolate genome. Corresponding positions in the reference sequence (GenBank: DQ217792) are shown between square brackets.

To further validate that MARV-ΔVP30-eGFP does not escape confinement to the transduced cell lines, both virus isolates were passaged an additional six times on Vero E6 cells transduced with codon-optimized MARV VP30. Supernatans from each passage was used to infect fresh transduced and untransduced Vero E6 cells. For each passage, >95% eGFP expression could be observed in all replicates of the transduced cells six days post-infection, while no eGFP expression or cytopathogenic effect was ever observed in untransduced cells, further confirming the biological confinement of these viruses.

### Screening of compound libraries for MARV

Following the characterization of our newly developed biologically contained MARV, we next exploited this system to screen for potential MARV inhibitors. Compounds, spotted in 96-well plates, were kindly provided to us by the Medicines for Malaria Venture, under the form of the Pandemic Response Box (400 compounds) and the Covid Box (160 compounds). All 560 compounds were initially screened in a 96-well format using four-fold dilutions ranging from 50 μM to 0.78 μM. Compound activity and toxicity were assessed by high-content imaging by scoring the declines in the fraction of eGFP-positive cells and the total number of cells, respectively. Additional compound in powder form was acquired for thirteen compounds that showed at least 40% more activity than toxicity in two or more of the tested dilutions. These compounds were then retested over a wider concentration range to determine their IC50 and CC50 values. Unfortunately, no compounds were found to strongly and selectively (selectivity indexes (SI) >7) inhibit MARV replication, except for the two known MARV inhibitors remdesivir and apilimod with (Figure 4). The activity of these two reference compounds was also assessed in the Huh-7-MARV-CO-VP30 cell line. In line with previously published data for these compounds, both showed increased toxicity in Huh-7 cells, with apilimod failing to show selective antiviral inhibition.

**Figure 4:**
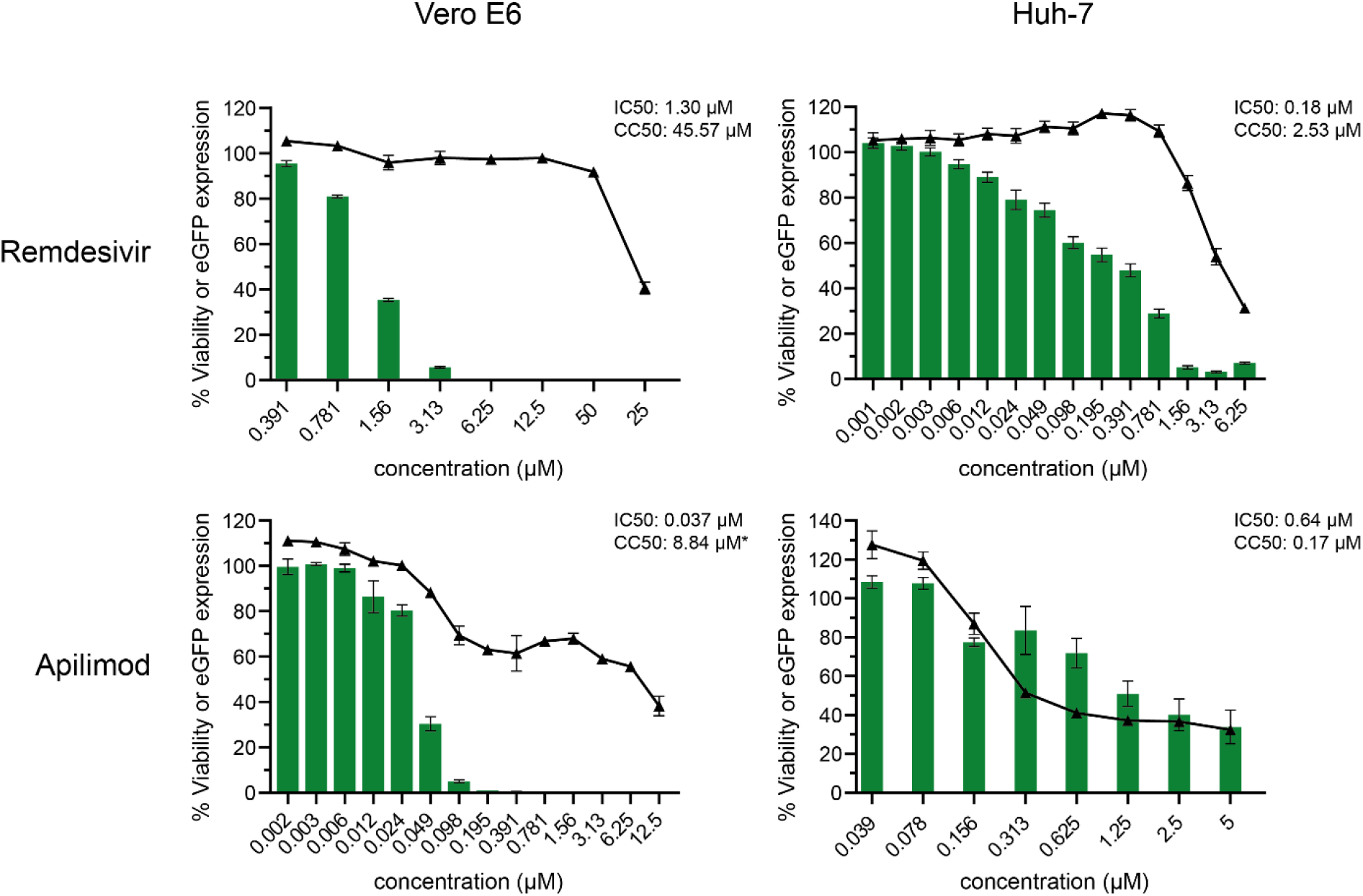
Anti-MARV activity and toxicity of remdesivir and apilimod. The graphs show the fraction of eGFP-expressing cells (green bars), or the total number of cells (black triangles), compared to the untreated control. Apilimod shows selective inhibition of MARV-ΔVP30-eGFP in Vero E6 cells, albeit with moderate toxicity, but not in Huh-7 cells. Conversely, remdesivir shows selective activity in both cell lines. IC50 and CC50 were calculated using a four-parameter non-linear regression model in GraphPad Prism v9.3.1. *For the CC50 value of apilimod in Vero E6 cells, an accurate curve could not be fitted, so the absolute CC50 value is shown. At least six replicates across two different plates were used for each condition. Error bars denote the standard error of mean.

### Adaptation to 384-well format

As a step-up for future work, we also verified the general applicability of our assay in 384-well plates by determining the minimal amount of virus that gives reliable infection of target cells. At a seeding density of 6,500 cells/well, a virus dilution series showed optimal virus growth at titers exceeding 96 PFU/well (~0.01 PFU/cell), with reproducibility and overall infectivity declining rapidly at further dilutions (Figure 5). 0.01 PFU/cell was therefore chosen as the optimal assay titer, yielding an average of 94.9% eGFP-positive cells (SD: 2.9%). Coupled with the near absence of false positivity in the negative control (average: <0.1%, SD: 0.2%), this results in a Z’-score of 0.9, confirming the usability of this format for high-throughput compound screening.

**Figure 5:**
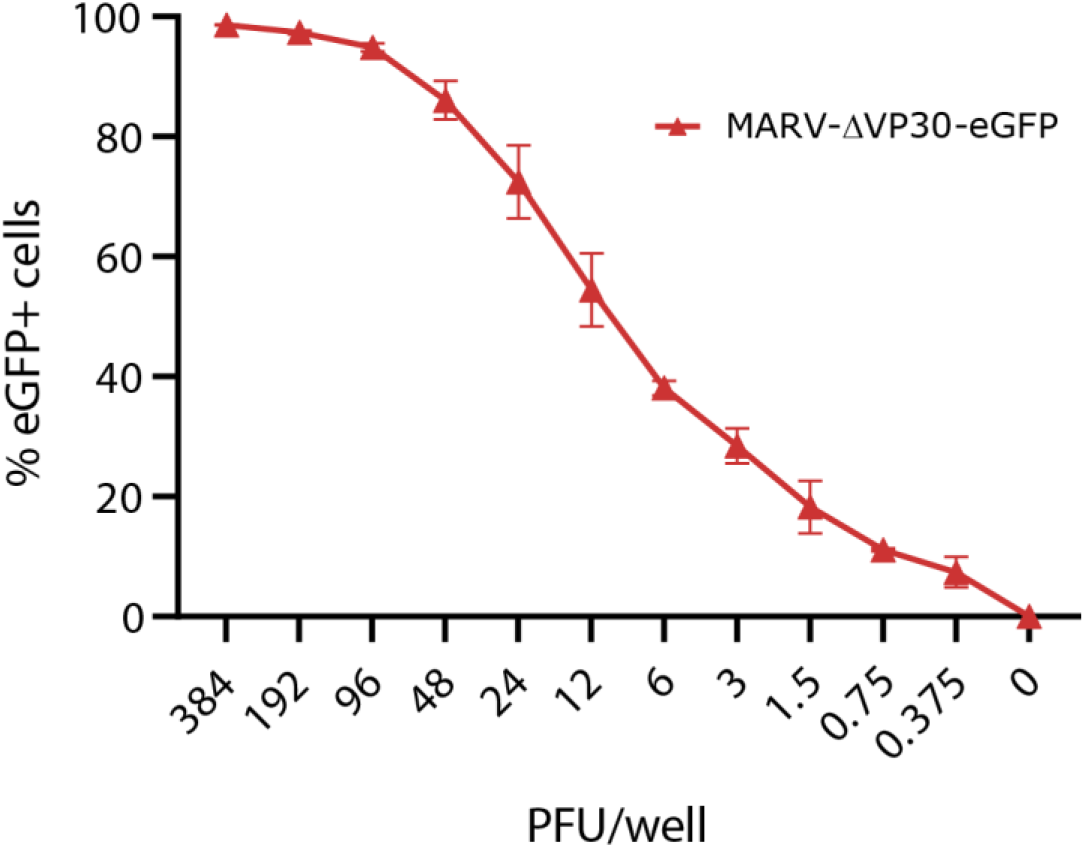
Minimal virus titer required for homogenous infection in 384-well plates. Vero E6-MARV-CO-VP30 cells were infected with MARV-ΔVP30-eGFP isolate B and the fraction of eGFP-positive cells was determined by high-content imaging six days post-infection. Different viral titers tested are expressed in plaque forming units (PFU)/well. Data from sixty-four replicates from two independent experiments. Error bars denote the standard error of mean.

## Discussion

In this study, we describe the rescue of biologically contained MARV and its confinement to stably transduced cell lines expressing VP30. Biological containment has previously been shown to be feasible for EBOV, and biologically confined EBOV has been used to evaluate the effect of several interferon-stimulated genes on EBOV replication, to study the role of receptor tyrosine kinases in Ebola virus entry and to screen for potential EBOV inhibitors [10, 13, 29, 30]. Conversely, for MARV, our study represents the first proof-of-concept that biological containment is possible. Only a limited number of MARV rescue systems have been developed so far and several research groups have noted difficulties in rescuing MARV, often requiring additional optimization of rescue conditions [8, 12, 26]. Also for the biologically contained MARV described here, rescue proved difficult to achieve. Because the replication of MARV-ΔVP30-eGFP is dependent on the presence of sufficient in trans-provided VP30 protein, rescue and subsequent propagation of MARV-ΔVP30-eGFP was unsuccessful in cell lines transduced with wild-type MARV VP30, which showed only limited protein expression. Codon optimization of the VP30 gene improved protein expression and in combination with codon optimization of the VP35, NP and L support plasmids yielded sufficient viral proteins for virus rescue. In line with observations by Enterlein et *al*. and Albariño et *al*., rescue using BSR-T7/5 cells, which are easy to transfect and show high protein expression levels after transduction, was hindered by the high turnover rate of these cells, presumably resulting in premature cell death before virus rescue could be achieved [12, 26]. To circumvent this issue, we performed the rescue directly in Vero E6 cells, and initial signs of virus production could be observed eight days post-transfection. It remains unclear why rescue takes such a long time to be achieved, unlike direct infection with rescued virus, which results in detectable eGFP expression within 24-48 hours.

While MARV-ΔVP30-eGFP growth was only efficient in Vero E6 cells expressing codon-optimized MARV VP30, some eGFP expression could also be observed in cells transduced with regular MARV or EBOV VP30. In the former case, eGFP expression was limited to a few small cell clusters per well. Likely, only a limited fraction (<0.1%) of the Vero E6-MARV-VP30 cells form sufficient VP30 protein to support virus transcription and replication. Only these cells and their progeny can efficiently replicate MARV-ΔVP30-eGFP, explaining the limited number and size of the observed eGFP-positive cell clusters. In cells transduced with EBOV VP30, infection with MARV-ΔVP30-eGFP results in widespread but weak eGFP expression. This is in line with the observations of Enterlein et al., who noted that rescue of recombinant MARV was possible when using EBOV VP30, although at reduced efficiency [12]. Interestingly, MARV-ΔVP30-eGFP isolate A, while behaving near identical to isolate B in other set-ups, seems to yield higher eGFP-expression levels when infecting Vero E6-EBOV-VP30 cells (Figure 3D). Although this increase in eGFP expression appears to be caused by an increased fraction of eGFP-expressing cells, close comparison of the images of the different isolates revealed that the amount of eGFP within cells was slightly higher for isolate A, resulting in more cells reaching the threshold to be counted as true positives. This slight increase in eGFP expression can potentially be explained by the addition of an extra adenosine residue in the polyadenylation signal of the eGFP gene, a mutation that has previously been described as a result of Vero E6-specific cell adaptation [31]. While experiments performed using bicistronic EBOV minigenomes have shown the impact of similar mutations to be limited, even a minor effect might be sufficient to explain the observed minimal difference between the two isolates [32].

For the generation of the cell lines to which the viruses are confined, lentiviral transduction was used. Compared to the co-transfection protocol previously used by Halfmann et al., lentiviral transduction yields a higher fraction of cells expressing both the gene of interest and the selection marker [10]. Furthermore, because the gene of interest and the selection marker are coupled with an IRES as part of the same transcriptional unit, it is ensured that all cells that survive antibiotic selection will also express the gene of interest. Even though IRES-coupled genes are known to generate uneven transcript levels, a negative impact of this unequal balance can be avoided by placing the resistance marker downstream of the IRES, as was done in our constructs [33]. Transduction using a bicistronic lentiviral vector also helps avoiding gene silencing, ultimately resulting in the stable expression of the gene of interest by the entire cell population [34]. By establishing stable cell lines that show uniform and reliable expression of VP30, both assay performance and reproducibility can be maximized.

The key advantage of the systems described here is that they can be used outside of a BSL-4 setting, thereby making research using replication-competent filoviruses available to a much bigger research community. Other virus alternatives that can be used in lower biosafety settings have previously been developed for EBOV and MARV, including minigenome systems, VLP systems and even trVLP systems [8, 9, 35, 36, 37]. While each of these surrogate systems has its advantages, they all have their limitations, including their dependency on repeated transfections or being limited to specific parts of the virus life cycle. Conversely, biologically contained viruses are actual replication-competent viruses that can be used to study all parts of the virus life cycle and that can be used in a variety of assays, although they may not be the optimal choice for studies on the role and function of VP30.

One application where this system can be particularly advantageous is the high-throughput screening of small-molecule compound libraries. Although some monoclonal antibody therapies have proven their benefit for the treatment of EVD, both EVD and MVD still have a poor prognosis due to limited treatment options [6]. Small-molecule compounds offer the advantage of being generally easy to develop and cheap to produce. Furthermore, millions of small-molecule compounds have already been developed, offering a potential source for the rapid identification of efficacious filovirus inhibitors. In recent years, screening of small-molecule libraries has already identified several promising lead compounds for the development of new filovirus antivirals [38]. However, because of the technical limitations of BSL-4 facilities, these compound screenings are typically performed using virus alternatives, the positive hits of which are not always directly applicable to wild-type virus. The biologically contained virus systems described here circumvent these limitations by being safe to handle in a BSL-2 setting while still providing biologically relevant results, thereby providing much lower false-positivity rates than sometimes observed in other BSL-2 assays. Additionally, the assay described here is already optimized to be used in combination with 384-well plates and can be fully automated. These optimizations enable the high-throughput screening of large (>100,000) compound libraries, making this system a highly interesting alternative for all currently developed assays. One remaining limitation is the dependency on specific transgene cells expressing MARV VP30, but, as demonstrated here, lentiviral transduction provides a robust method for the creation of such cell lines. We already showed the efficient implementation of two different VP30^+^ cell lines. Using this method, we are currently developing and optimizing complementary assays using alternative cell lines, to overcome some of the shortcomings associated with working with Vero E6 and Huh-7 cells.

In conclusion, we show that, comparable to EBOV, it is feasible to confine MARV to a specific cell line, allowing its handling at lower biosafety levels. Furthermore, we describe the rescue of MARV-ΔVP30-eGFP and the generation of cell lines to which it is confined via lentiviral transduction with MARV VP30. Lastly, we also used this system to screen the MMV Pandemic Response Box and Covid Box compound collections and illustrated its potential to be used in 384-well format, highlighting its usability for (high-throughput) compound screening assays.

## Author contributions

This study was conceived by BV and PM. Experimental work was performed by BV and JSt. High-content imaging was performed by WC and JSc. Data analysis was performed by BV. PM, LN and KV supplied reagents and materials. BV and PM drafted the manuscript. All authors read and approved the final version of the manuscript.

## Acknowledgments

The authors wish to thank the Medicines for Malaria Venture for providing compounds, prof. R. Bartenschlager, University of Heidelberg, Germany for providing the Huh-7 cells, dr. K. Conzelmann, Friedrich-Loeffler-Institut, Germany for providing the BST7/5 cells and prof. S. Becker, Philipps-Universität, Marburg, Germany for providing the T7-3M-Luc-5M minigenome and MARV support plasmids. The authors also wish to thank dr. Dirk Jochmans and Bram Van Holm for the insightful discussions. B.V. was supported by a FWO SB grant for strategic basic research of the “Fonds Wetenschappelijk Onderzoek”/Research foundation Flanders (1S28617N). Part of this research work was performed using the ‘Caps-It’ research infrastructure (project ZW13-02) that was financially supported by the Hercules Foundation (FWO) and Rega Foundation, KU Leuven.

## Supplementary Data

**Supplementary Table S1:**
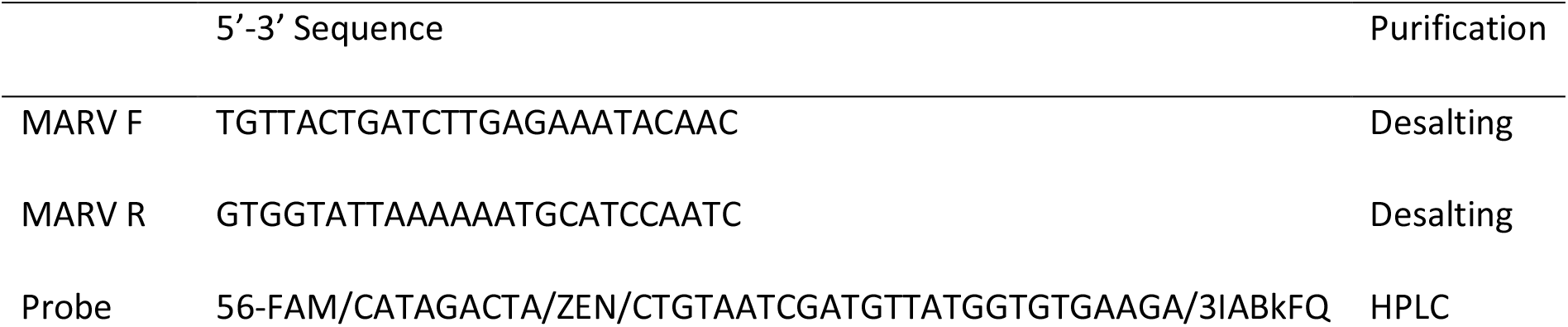
qPCR primers and probes

Supplementary Figure S1: Western Blot of VP30-expressing cell lines

Supplementary Figure S2: Growth kinetics and biological containment of MARV-ΔVP30-eGFP

Supplementary Figure S3: Growth kinetics of MARV-ΔVP30-eGFP eGFP in cells expressing wild-type MARV or EBOV VP30

## Supplementary Figure Legends

**Supplementary Figure S1:**
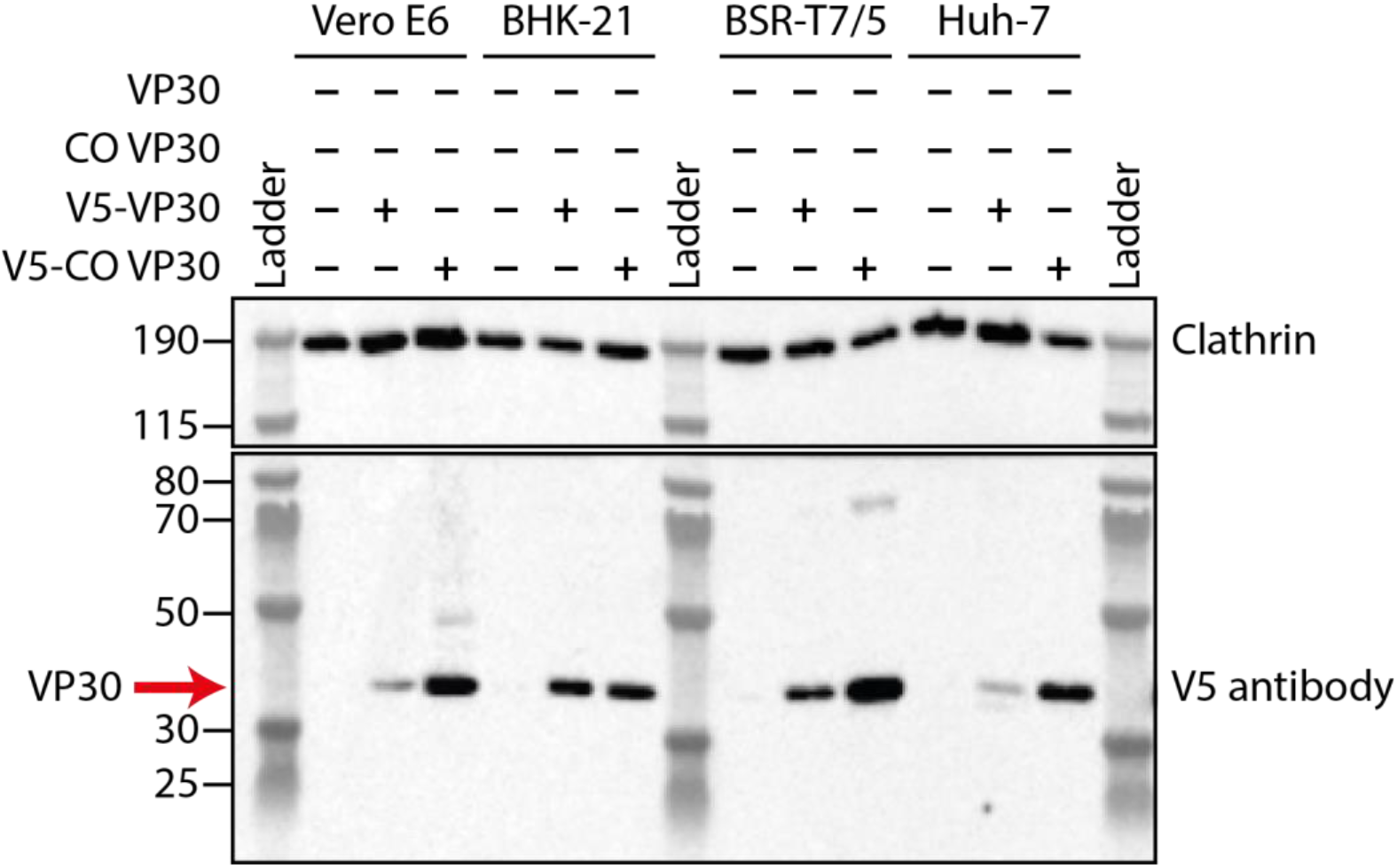
Western blot illustrating VP30 expression after lentiviral transduction. Clathrin was used as a loading control for all lanes. Blotting using a V5-specific antibody shows an increased expression of V5-tagged MARV VP30 after codon optimization, particularly in Vero E6 and Huh-7 cells. Blot exposure time: 100s. CO = codon optimized.

**Supplementary Figure S2:**
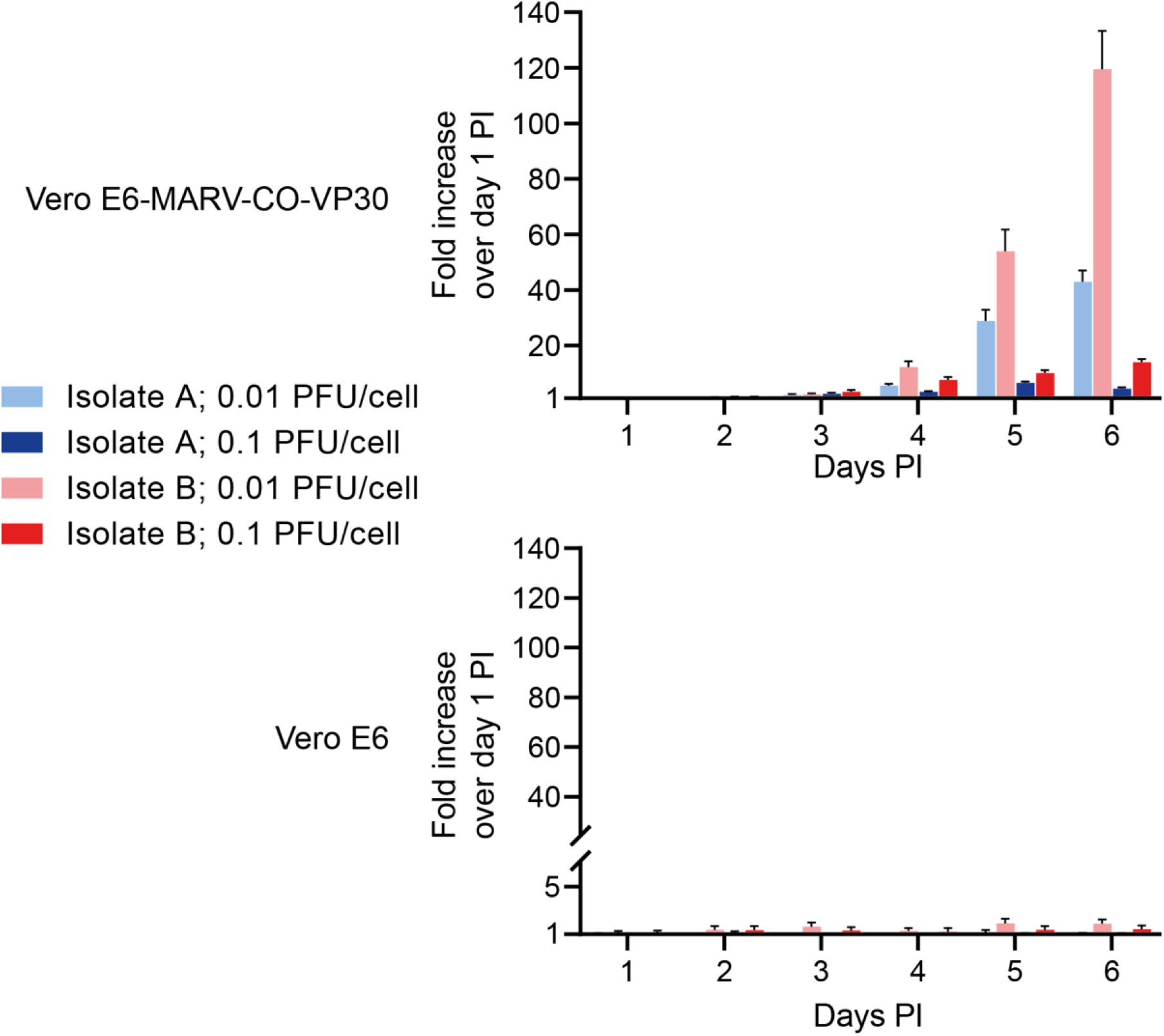
Graphs showing the ratio of genome copies compared to the amount at day 1 post-infection (PI), as measured by qPCR. At least three replicates are included for each condition. Error bars denote standard error.

**Supplementary Figure S3:**
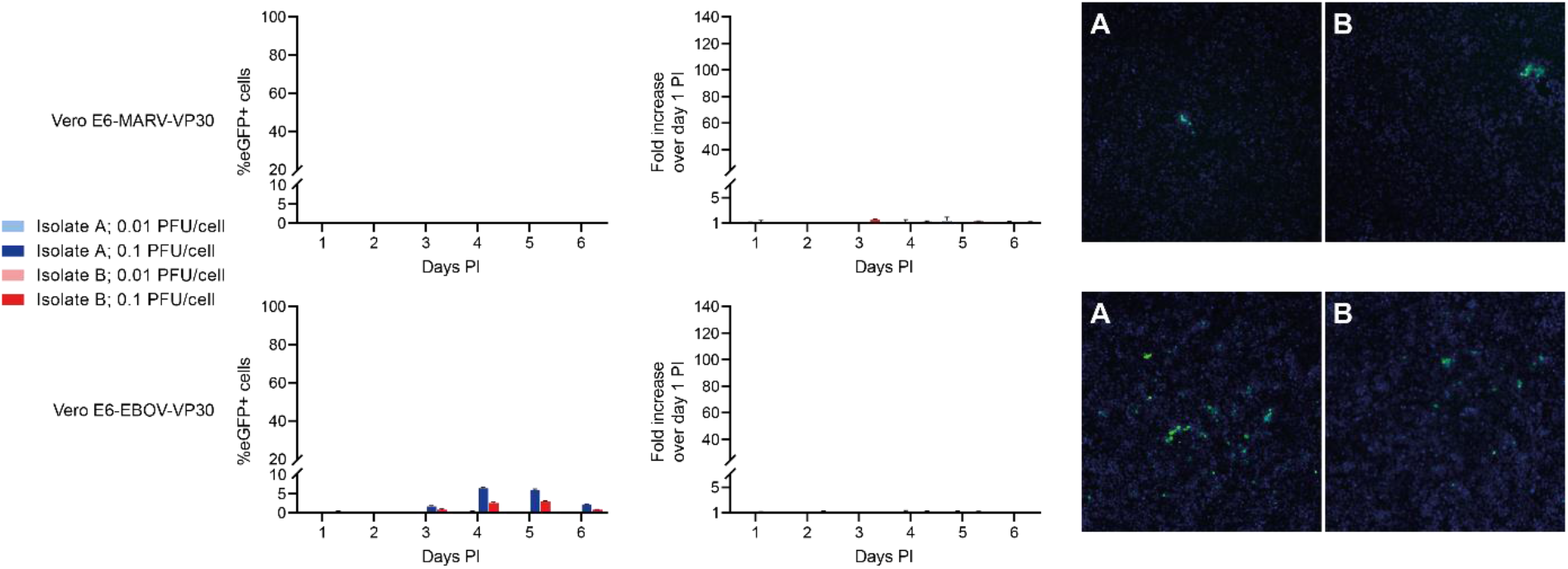
The left graph shows the background-corrected fraction of eGFP-positive cells, measured by high-content imaging. The middle graphs show the ratio of genome copies compared to the amount at day 1 post-infection (PI), as measured by qPCR. The right column shows a representative image of a well infected with 0.1 PFU/well, five days post-infection. Green cells express eGFP and all cells are nuclear stained with Hoechst 33342 (blue). PFU = plaque forming units. At least three replicates are included for each condition. Error bars denote standard error.

## References

1. Kuhn JH, Amarasinghe GK, Basler CF, et al. ICTV Virus Taxonomy Profile: Filoviridae. The Journal of general virology. 2019 Jun;100(6):911–912. doi: 10.1099/jgv.0.001252. PubMed PMID: 31021739; PubMed Central PMCID: PMC7011696.

2. Languon S, Quaye O. Filovirus Disease Outbreaks: A Chronological Overview. Virology: research and treatment. 2019;10:1178122X19849927. doi: 10.1177/1178122X19849927. PubMed PMID: 31258326; PubMed Central PMCID: PMC6589952.

3. U.S. Food and Drug Administration. First FDA-approved vaccine for the prevention of Ebola virus disease, marking a critical milestone in public health preparedness and response, 2019. [Accessed on May 19th, 2021]. Available from: https://www.fda.gov/news-events/press-announcements/first-fda-approved-vaccine-prevention-ebola-virus-disease-marking-critical-milestone-public-health

4. Henao-Restrepo AM, Camacho A, Longini IM, et al. Efficacy and effectiveness of an rVSV-vectored vaccine in preventing Ebola virus disease: final results from the Guinea ring vaccination, open-label, cluster-randomised trial (Ebola Ca Suffit!). Lancet. 2017 Feb 4;389(10068):505–518. doi: 10.1016/S0140-6736(16)32621-6. PubMed PMID: 28017403; PubMed Central PMCID: PMC5364328.

5. European Medicines Agency. New vaccine for prevention of Ebola virus disease recommended for approval in the European Union, 2020. [Accessed on August 31th, 2021]. Available from: https://www.ema.europa.eu/en/news/new-vaccine-prevention-ebola-virus-disease-recommended-approval-european-union

6. Mulangu S, Dodd LE, Davey RT, Jr., et al. A Randomized, Controlled Trial of Ebola Virus Disease Therapeutics. The New England journal of medicine. 2019 Dec 12;381(24):2293–2303. doi: 10.1056/NEJMoa1910993. PubMed PMID: 31774950.

7. Chosewood LC, Wilson DE, Centers for Disease Control and Prevention (U.S.), et al. Biosafety in microbiological and biomedical laboratories. 5th ed. Washington, D.C.: U.S. Dept. of Health and Human Services, Public Health Service, Centers for Disease Control and Prevention, National Institutes of Health; 2009. (HHS publication; no (CDC) 21-1112).

8. Schmidt KM, Muhlberger E. Marburg Virus Reverse Genetics Systems. Viruses. 2016 Jun 22;8(6). doi: 10.3390/v8060178. PubMed PMID: 27338448; PubMed Central PMCID: PMC4926198.

9. Hoenen T, Groseth A, de Kok-Mercado F, et al. Minigenomes, transcription and replication competent virus-like particles and beyond: reverse genetics systems for filoviruses and other negative stranded hemorrhagic fever viruses. Antiviral research. 2011 Aug;91(2):195–208. doi: 10.1016/j.antiviral.2011.06.003. PubMed PMID: 21699921; PubMed Central PMCID: PMC3586226.

10. Halfmann P, Kim JH, Ebihara H, et al. Generation of biologically contained Ebola viruses. Proceedings of the National Academy of Sciences of the United States of America. 2008 Jan 29;105(4):1129–33. doi: 10.1073/pnas.0708057105. PubMed PMID: 18212124; PubMed Central PMCID: PMC2234103.

11. Bach S, Demper JC, Grunweller A, et al. Regulation of VP30-Dependent Transcription by RNA Sequence and Structure in the Genomic Ebola Virus Promoter. Journal of virology. 2020 Dec 2. doi: 10.1128/JVI.02215-20. PubMed PMID: 33268520; PubMed Central PMCID: PMC8092829.

12. Enterlein S, Volchkov V, Weik M, et al. Rescue of recombinant Marburg virus from cDNA is dependent on nucleocapsid protein VP30. Journal of virology. 2006 Jan;80(2):1038–43. doi: 10.1128/JVI.80.2.1038-1043.2006. PubMed PMID: 16379005; PubMed Central PMCID: PMC1346851.

13. Vanmechelen B, Stroobants J, Chiu W, et al. Identification of novel Ebola virus inhibitors using biologically contained virus. Antiviral research. 2022 Apr;200:105294. doi: 10.1016/j.antiviral.2022.105294. PubMed PMID: 35337896.

14. Medicines for Malaria Venture. The COVID Box, 2021. [Accessed on June 16th, 2021]. Available from: https://www.mmv.org/mmv-open/covid-box

15. Medicines for Malaria Venture. The Pandemic Response Box, 2021. [Accessed on June 16th, 2021]. Available from: https://www.mmv.org/mmv-open/pandemic-response-box

16. Nelson EA, Dyall J, Hoenen T, et al. The phosphatidylinositol-3-phosphate 5-kinase inhibitor apilimod blocks filoviral entry and infection. PLoS neglected tropical diseases. 2017 Apr;11(4):e0005540. doi: 10.1371/journal.pntd.0005540. PubMed PMID: 28403145; PubMed Central PMCID: PMC5402990.

17. Lo MK, Jordan R, Arvey A, et al. GS-5734 and its parent nucleoside analog inhibit Filo-, Pneumo-, and Paramyxoviruses. Scientific reports. 2017 Mar 6;7:43395. doi: 10.1038/srep43395. PubMed PMID: 28262699; PubMed Central PMCID: PMC5338263.

18. Buchholz UJ, Finke S, Conzelmann KK. Generation of bovine respiratory syncytial virus (BRSV) from cDNA: BRSV NS2 is not essential for virus replication in tissue culture, and the human RSV leader region acts as a functional BRSV genome promoter. Journal of virology. 1999 Jan;73(1):251–9. doi: 10.1128/JVI.73.1.251-259.1999. PubMed PMID: 9847328; PubMed Central PMCID: PMC103829.

19. Wenigenrath J, Kolesnikova L, Hoenen T, et al. Establishment and application of an infectious virus-like particle system for Marburg virus. The Journal of general virology. 2010 May;91(Pt 5):1325–34. doi: 10.1099/vir.0.018226-0. PubMed PMID: 20071483.

20. Goya S, Valinotto LE, Tittarelli E, et al. An optimized methodology for whole genome sequencing of RNA respiratory viruses from nasopharyngeal aspirates. PloS one. 2018;13(6):e0199714. doi: 10.1371/journal.pone.0199714. PubMed PMID: 29940028; PubMed Central PMCID: PMC6016902.

21. Li H. Minimap2: pairwise alignment for nucleotide sequences. Bioinformatics. 2018 Sep 15;34(18):3094–3100. doi: 10.1093/bioinformatics/bty191. PubMed PMID: 29750242; PubMed Central PMCID: PMC6137996.

22. Vanmechelen B, Stroobants J, Vermeire K, et al. Advancing Marburg virus antiviral screening: Optimization of a novel T7 polymerase-independent minigenome system. Antiviral research. 2021 Jan;185:104977. doi: 10.1016/j.antiviral.2020.104977. PubMed PMID: 33220335.

23. Koehler A, Kolesnikova L, Welzel U, et al. A Single Amino Acid Change in the Marburg Virus Matrix Protein VP40 Provides a Replicative Advantage in a Species-Specific Manner. Journal of virology. 2016 Feb 1;90(3):1444–54. doi: 10.1128/jvi.02670-15. PubMed PMID: 26581998; PubMed Central PMCID: PMCPMC4719610. eng.

24. Albariño CG, Wiggleton Guerrero L, Lo MK, et al. Development of a reverse genetics system to generate a recombinant Ebola virus Makona expressing a green fluorescent protein. Virology. 2015 2015/10/01/;484:259–264. doi: https://doi.org/10.1016/j.virol.2015.06.013.

25. Dolnik O, Kolesnikova L, Welsch S, et al. Interaction with Tsg101 is necessary for the efficient transport and release of nucleocapsids in marburg virus-infected cells. PLoS pathogens. 2014 Oct;10(10):e1004463. doi: 10.1371/journal.ppat.1004463. PubMed PMID: 25330247; PubMed Central PMCID: PMCPMC4199773. eng.

26. Albarino CG, Uebelhoer LS, Vincent JP, et al. Development of a reverse genetics system to generate recombinant Marburg virus derived from a bat isolate. Virology. 2013 Nov;446(1-2):230–7. doi: 10.1016/j.virol.2013.07.038. PubMed PMID: 24074586; PubMed Central PMCID: PMC5683708.

27. Neumann G, Feldmann H, Watanabe S, et al. Reverse Genetics Demonstrates that Proteolytic Processing of the Ebola Virus Glycoprotein Is Not Essential for Replication in Cell Culture. 2002;76(1):406–410. doi: doi:10.1128/JVI.76.1.406-410.2002.

28. Volchkov VE, Volchkova VA, Mühlberger E, et al. Recovery of Infectious Ebola Virus from Complementary DNA: RNA Editing of the GP Gene and Viral Cytotoxicity. 2001;291(5510):1965–1969. doi: doi:10.1126/science.1057269.

29. Kuroda M, Halfmann PJ, Hill-Batorski L, et al. Identification of interferon-stimulated genes that attenuate Ebola virus infection. Nature communications. 2020 Jun 11;11(1):2953. doi: 10.1038/s41467-020-16768-7. PubMed PMID: 32528005; PubMed Central PMCID: PMC7289892.

30. Kuroda M, Halfmann P, Kawaoka Y. HER2-mediated enhancement of Ebola virus entry. PLoS pathogens. 2020 Oct;16(10):e1008900. doi: 10.1371/journal.ppat.1008900. PubMed PMID: 33052961; PubMed Central PMCID: PMC7556532.

31. Tsuda Y, Hoenen T, Banadyga L, et al. An Improved Reverse Genetics System to Overcome Cell-Type-Dependent Ebola Virus Genome Plasticity. The Journal of infectious diseases. 2015 Oct 1;212 Suppl 2:S129–37. doi: 10.1093/infdis/jiu681. PubMed PMID: 25810440; PubMed Central PMCID: PMC4564527.

32. Brauburger K, Boehmann Y, Krahling V, et al. Transcriptional Regulation in Ebola Virus: Effects of Gene Border Structure and Regulatory Elements on Gene Expression and Polymerase Scanning Behavior. Journal of virology. 2016 Feb 15;90(4):1898–909. doi: 10.1128/JVI.02341-15. PubMed PMID: 26656691; PubMed Central PMCID: PMC4733972.

33. Osti D, Marras E, Ceriani I, et al. Comparative analysis of molecular strategies attenuating positional effects in lentiviral vectors carrying multiple genes. Journal of virological methods. 2006 Sep;136(1-2):93–101. doi: 10.1016/j.jviromet.2006.04.003. PubMed PMID: 16690138.

34. Wiznerowicz M, Trono D. Harnessing HIV for therapy, basic research and biotechnology. Trends in biotechnology. 2005 Jan;23(1):42–7. doi: 10.1016/j.tibtech.2004.11.001. PubMed PMID: 15629857.

35. Hoenen T, Watt A, Mora A, et al. Modeling the lifecycle of Ebola virus under biosafety level 2 conditions with virus-like particles containing tetracistronic minigenomes. Journal of visualized experiments: JoVE. 2014 Sep 27(91):52381. doi: 10.3791/52381. PubMed PMID: 25285674; PubMed Central PMCID: PMC4828136.

36. Gan T, Zhou D, Huang Y, et al. Development of a New Reverse Genetics System for Ebola Virus. mSphere. 2021 May 5;6(3). doi: 10.1128/mSphere.00235-21. PubMed PMID: 33952663; PubMed Central PMCID: PMCPMC8103987. eng.

37. Tao W, Gan T, Guo M, et al. Novel Stable Ebola Virus Minigenome Replicon Reveals Remarkable Stability of the Viral Genome. Journal of virology. 2017 Nov 15;91(22). doi: 10.1128/jvi.01316-17. PubMed PMID: 28878087; PubMed Central PMCID: PMCPMC5660472. eng.

38. Edwards MR, Basler CF. Current status of small molecule drug development for Ebola virus and other filoviruses. Current opinion in virology. 2019 Apr;35:42–56. doi: 10.1016/j.coviro.2019.03.001. PubMed PMID: 31003196; PubMed Central PMCID: PMC6556423.

